# Multi-scale modeling shows that dielectric differences make Na_V_ channels faster than K_V_ channels

**DOI:** 10.1101/2020.05.11.088559

**Authors:** Luigi Catacuzzeno, Luigi Sforna, Fabio Franciolini, Robert S. Eisenberg

**Author notes:** **Correspondence to:** Luigi Catacuzzeno, Department of Chemistry, Biology and Biotechnology, University of Perugia, via Elce di Sotto 8, 06123 Perugia - Italy, **Email address:**.

## Abstract

The generation of action potentials in excitable cells requires different activation kinetics of voltage gated Na (Na_V_) and K (K_V_) channels. Na_V_ channels activate much faster and allow the initial Na^+^ influx that generates the depolarizing phase and propagates the signal. Recent experimental results suggest that the molecular basis for this kinetic difference is an amino acid side chain located in the gating pore of the voltage sensor domain, which is a highly conserved isoleucine in K_V_ channels, but an equally highly conserved threonine in Na_V_ channels. Mutagenesis suggests that the hydrophobicity of this side chain in Shaker K_V_ channels regulates the energetic barrier that gating charges need to overcome to move through the gating pore, and ultimately the rate of channel opening. We use a multi-scale modeling approach to test this hypothesis. We use high resolution molecular dynamics to study the effect of the mutation on polarization charge within the gating pore. We then incorporate these results in a lower resolution model of voltage gating to predict the effect of the mutation on the movement of gating charges. The predictions of our hierarchical model are fully consistent with the tested hypothesis, thus suggesting that the faster activation kinetics of Na_V_ channels comes from a stronger dielectric polarization by threonine (Na_V_ channel) produced as the first gating charge enters the gating pore, compared to isoleucine (K_V_ channel).

**eTOC Summary:** Voltage-gated Na^+^ channels activate faster than K^+^ channels in excitable cells. Catacuzzeno et al. develop a model that shows how the dielectric properties of a divergent side-chain produce this difference in speed.

## Introduction

The most important role of voltage-gated sodium (Nav) and potassium (Kv) channels is their ability to generate action potentials, all-or-none (i.e., binary) rapid and transient changes of the electric potential difference across the plasma membrane, that propagate signals long distances in the nervous system. This ability is essentially due to the different timing with which these two channels respond to a potential change (Figure 1A): Nav channels activate rapidly upon an initial depolarization of the membrane, allowing a Na current to enter the cell and induce a further membrane depolarization along the nerve during the first millisecond or so after the initial trigger. By contrast Kv channels remain essentially closed during this initial interval, and open only later, causing the repolarization of the membrane to the resting condition (Bezanilla, Rojas, and Taylor 1970; Hodgkin and Huxley 1952; Rojas, Bezanilla, and Taylor 1970).

**Figure 1.**
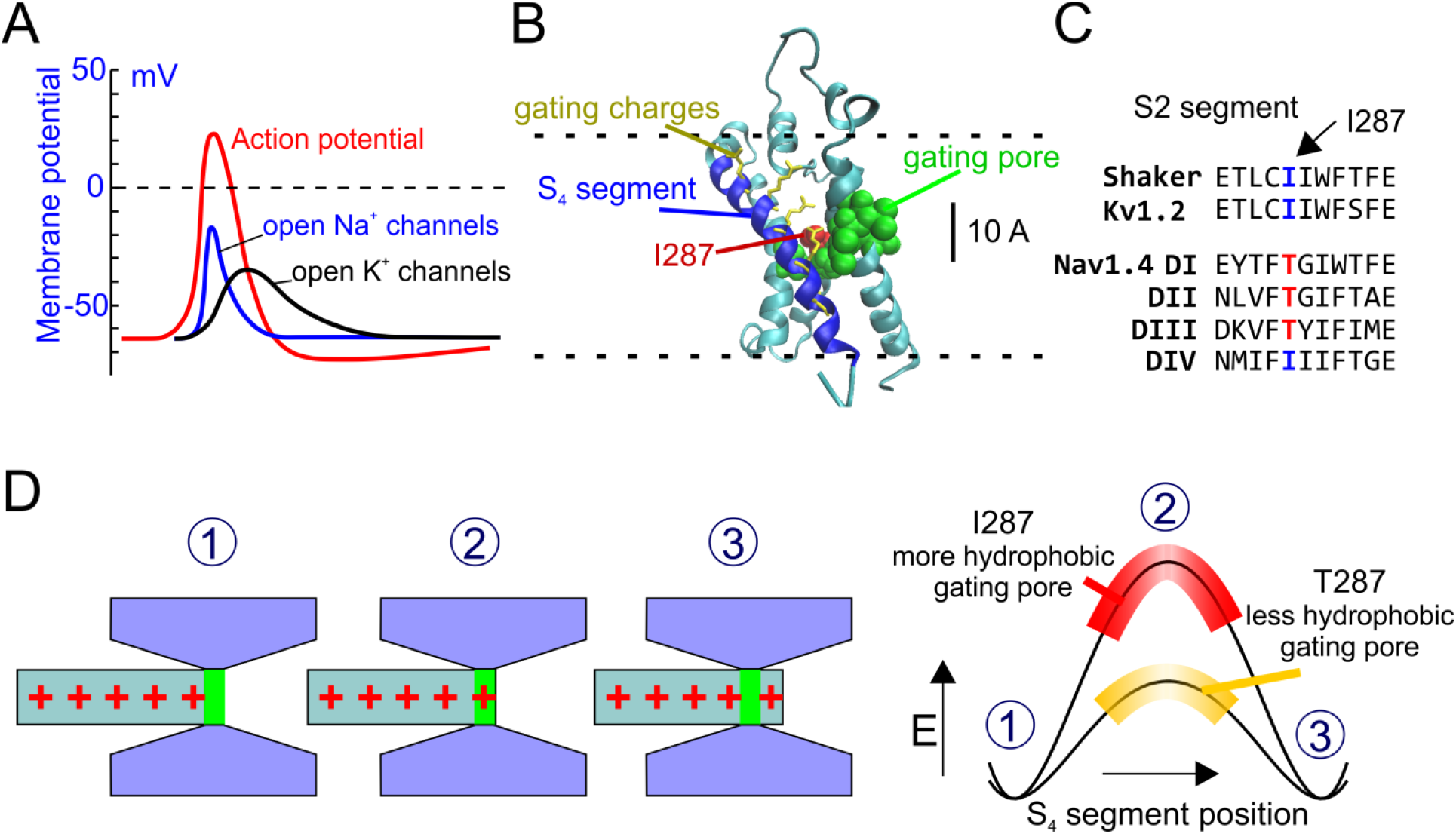
**A)** Schematic plot showing the activity of Nav and Kv channels during an action potential (red line). Nav channels open more rapidly than Kv channels, creating the depolarizing phase of the action potential. **B)** 3D structure of the active state of the VSD of the Shaker channel from (Henrion et al. 2012). The S4 segment is in blue and the S1-S3 segments are in cyan. The hydrophobic residues forming the gating pore (V236, I237, S240, I241, F244, C286, I287, F290, A319, I320;(Lacroix et al. 2014)) are shown in van der Waals (VdW) representation (green). The residue 287 is colored in red. Gating charge side chains are shown in licorice representation and colored in yellow. **C)** Portion of the S2 segments of several voltage sensors belonging to Shaker, Kv and fast Nav channels, showing the highly conserved isoleucine at position 287 in Shaker-like potassium channels, and the threonine residue at the corresponding position in the fast Nav1.4 channel. **D)** Schematic representation of the hypothesis tested in this paper. Left: the S4 voltage sensor is represented as a cylinder with the gating charges in red. The rest of the VSD is in gray, and forms the gating pore and the intra- and extracellular vestibules. Right: energetic profiles encountered by the first gating charge across the gating pore, in presence of two different hydrophobicity levels of the gating pore. When the gating charge enters the gating pore there is an increase in the energy of the system, due to the destabilizing (hydrophobic) environment within the gating pore. The more hydrophobic is the gating pore, the higher will be this energy barrier and slower the activation rate.

The molecular and physical basis for the different kinetics of Nav and Kv channels has long been sought. The recently determined crystal structure of the voltage sensor domain (VSD) of Kv and Nav channels has allowed investigation of the structure-function relationship of the voltage sensor, its environment, and the forces controlling its movement. In both channels the VSD is formed by the transmembrane segments 1-4 (S1-S4), with the S4 segment containing positively charged residues every third amino acid, and forming the voltage sensor, the structure that senses the electric field applied across the membrane and moves upon depolarization (Long et al. 2007; Long, Campbell, and MacKinnon 2005; Pan et al. 2018; Payandeh et al. 2011).

The S4 segment is surrounded for most of its length by wide intracellular and extracellular vestibules where water and ions can easily access and balance the gating charges (Figure 1B). The two vestibules are separated by a series of mostly non polar residues that form a region, the gating pore, thought to be hardly accessible to water (green and red residues in Figure 1B). Several mutagenesis experiments suggest that some of these residues are in direct contact with the moving gating charges, and modulate the voltage-dependent properties of the channels (Lacroix et al. 2012, 2013, 2014; Lacroix and Bezanilla 2011; Tao et al. 2010).

One of the residues of the gating pore that determine the slow kinetics of Kv channels, compared to faster Nav channels, is isoleucine at position 287, located in the middle of the S2 segment (red in Figure 1B; notice that in this paper we will always use the residue numbering of the Shaker channel). This non polar, highly hydrophobic residue is found in many Kv channels. It is conserved by evolution presumably because of its essential role. By contrast, in the S2 segment of the first three domains of fast Nav channels, at corresponding position of isoleucine in Kv channels we consistently find the uncharged but much more hydrophilic residue threonine (Figure 1C; see also supplementary Figure 1 for a more extended sequence alignment) (Lacroix et al. 2013). This seems to be an example of divergent side-chain evolution, in which two related proteins (Nav and Kv channels) have selected two different side-chains as an adaptation to create different dielectric polarization and different activation kinetics, and thus to create a propagating action potential. Notably, when isoleucine at position 287 is mutated to threonine (or to other polar amino acids), the Shaker gating currents become much faster and similar to the activation rate of Nav channels gating currents (Lacroix et al. 2013, 2014). These results strongly suggest that the physical structure and the underlying mechanism controlling the activation rate of voltage-gated ion channels involve the hydrophobic strength, i.e., the polarizability of the gating pore, especially of the residue at position 287. They suggest that a highly non polar, hydrophobic gating pore creates an energetically unfavorable condition, a high energetic barrier (mostly a dielectric barrier) for the gating charges to move through upon depolarization, resulting in a slowing of the voltage sensor activation (Figure 1D). In this view, increasing the polarizability of this region by replacing a highly non polar side chain residue with a more polar amino acid lowers the dielectric energy barrier, provides counter charge (in the wall of the gating pore) for the gating charges, and ultimately increases the activation rate of the voltage sensor.

We test this mechanistic hypothesis using a multi-scale modeling approach; that is, an approach that investigates the system at two different time and space scales, in a series of hierarchical computational methods whereby the calculated quantities from a computational simulation at an atomistic scale are used to define the parameters of the model at a coarser, macroscopic scale, that satisfies conservations laws (Eisenberg 2018). In the specific case studied here, of voltage dependent gating, the idea translates into using Brownian modeling to macroscopically describe the system and obtain gating currents, and then using molecular dynamics (MD) simulations to determine the needed free parameters present in the Brownian model. It is difficult to determine these parameters experimentally but there is hope from the exciting work of the Boxer group (Fried, Bagchi, and Boxer 2014; Fried and Boxer 2017; Suydam 2006). The multi-scale approach appears feasible because assessing free parameters through MD simulations is much less susceptible to artifact and is less computationally demanding than simulating the overall movement of the voltage sensor. It also has the advantage of using a Brownian model that automatically satisfyes conservation laws of mass, current, and charge, and Poisson’s equations. It is difficult to satisfy Poisson’s equations, and conservation of current when using the periodic boundary conditions common in molecular dynamics.

## Methods

### Molecular dynamics simulations

The voltage sensor domain used in the simulation was taken from the Shaker model of Henrion et al. (2012), in the activated state. For MD simulations, the protein was embedded on a membrane environment made of POPC, with dimensions of 80×80 Å, and solvated in two steps. The distance between the maximum and minimum z coordinates and the water box edges in the z-axis was set to 15 Å. The system was first solvated below the membrane plane with a water box of dimensions 83.46, 80.05, 16.19 (x, y, z in Å) where 9462 TIP3 water molecules (Jorgensen et al. 1983) were placed. Then the system was solvated above the membrane plane with a water box of the dimensions 83.46, 80.05, 20.02 (x, y, z in Å) where a total of 9462 TIP3 water molecules were placed. After the addition of the water molecules, the system presented a total net charge of −2. The system was balanced and a total of 27 Cl ions plus 29 Na ions were added to the system making up a salt concentration of 0.15 mol/L. The MD simulations in the present study were performed employing the NAMD molecular dynamics package (Phillips et al. 2005). The CHARMM36 force field (Best et al. 2012; MacKerell et al. 1998) was used in all MD simulations. A distance cut-off of 12.0 Å was applied to short-range, non-bonded interactions, and of 10.0 Å for the smothering functions. Long-range electrostatic interactions were treated using the particle-mesh Ewald (PME) (Darden, York, and Pedersen 1993) method. Before the MD simulations all the systems were submitted to an energy minimization protocol for 2000 steps. After minimization an annealing of 0.29 ns was performed, with a temperature ramp of 0.24 ns of simulation where the temperature was raised from 60 K to 300 K. The pressure was maintained at 1 atm using Nosé-Hoover Langevin piston (Feller et al. 1995; Martyna, Tobias, and Klein 1994). The equations of motion were integrated using the r-RESPA multiple time step scheme (Phillips et al. 2005) to update the short-range interactions every 1 step and long-range electrostatics interactions every 2 steps. The time step of integration was chosen to be 2 fs for all simulations. During the Annealing the motion of the backbone of the protein was restrained. After the annealing a 10 ns equilibration was performed, while maintaining the restriction of the backbone motion, during which the temperature was maintained at 300 K using Langevin dynamics. The pressure was maintained at 1 atm using Nosé-Hoover Langevin piston. A distance cut-off of 12.0 Å was applied to short-range, non-bonded interactions, and 10.0 Å for the smothering functions. Long-range electrostatic interactions were treated using the particle-mesh Ewald (PME) method. Finally, we performed unrestricted MD simulations by applying different values of the electric field in the z direction. These simulations were used to assess the running average of total dipole moment, used to estimate the dielectric constant as described in the “Results” section.

### The Brownian model of voltage gating

A full description of the Brownian model of voltage gating can be found in (Catacuzzeno and Franciolini 2019; Catacuzzeno, Sforna, and Franciolini 2020). Briefly, in our model the VSD was approximated to a hourglass-shaped geometrical structure consisting of a cylindrical gating pore, having a length of 7 Å and a diameter of 10 Å, flanked by internal and external water accessible vestibules having a length of 3.15 nm each and a conical shape opening with a half angle of 15° into two hemispherical subdomains of bath solution, both having radius of 1 μm. The S4 charge profile (Z_S4_) was built by considering the six positive charges, whose mean distance between the charged atoms was determined from a 3D Shaker channel model, and each giving rise to a charge profile normally distributed with a standard deviation of 0.1 nm. The permanently charged profile (Z_F_), which in this case deserves the name fixed because it does not move, was similarly built by considering the vertical dimension of the C_α_ of all the charged residues on segments S1-S3 of the VSD.

Ions moved by electro-diffusion governed by the following flux conservation equation:

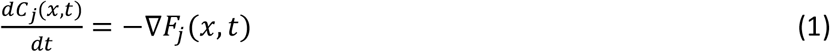

where *C_j_*(*x, t*) is the concentration of ion j, t is the time, ∇ is the spatial gradient operator, and *F_j_*(*x, t*) is the flux (mole per second per unit area) of ion j, given by the Nernst-Planck equation:

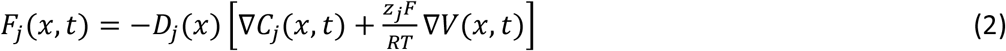

where *D_j_*(*x*) and *z_j_* are the diffusion coefficient profile and the valence of ion j, respectively, F, R and T have their usual meanings, and *V*(*x,t*) is the electrical voltage profile. Because ions move much faster in the bath than the voltage sensor moves in the gating a quasi-equilibrium assumption was made for ions electrodiffusion by imposing *F_j_*(*x, t*) = 0.

The S4 segment was assumed to move in one dimension as a Brownian particle, whose dynamics is governed by the following Langevin’s equation

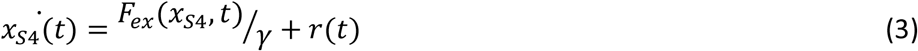

Here *x*_*S*4_(*t*) represents the position of the voltage sensor (distance between the R2-R3 midpoint and the center of the gating pore), *m* is the mass of the particle, *F_ex_*(*x*_S4_, *t*) is the external force acting on the particle, and R(t) is a random force due to the collision of the fluid and the rest of the protein on the S4 segment, which has a probability distribution with zero mean and second moment given by < *R*(*t*) *R*(*′*) >= 2 *k_B_T δ*(*t* – *t′*) where *k_B_* is the Boltzmann constant and *δ* is the delta function. The external force was assumed to have an electrical component, assessed from the electric voltage profile, and a spring component with force constant of 0.005 N/m to account for the force exerted by the channel permeation pore, which in this model has not been considered (Catacuzzeno and Franciolini 2019). *γ*, the friction coefficient of the S4 voltage-sensor, was set to 2*10^-6^ Kg/s based on the comparison between experimental and predicted macroscopic gating currents (notice that in Catacuzzeno et al., 2019 this parameter was set to twice this value). The dynamics of the S4 segment may also be described in terms of the time evolution of the probability density function profile, given by the following Fokker-Planck (FP) equation:

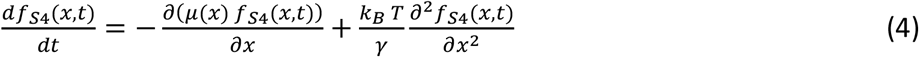

Here *f*_*S*4_ represents the probability density function for the position of the S4 segment and *μ*(*x*) is *F_ex_*(*x*_*S*4_, *t*)/*γ*. This partial differential equation was solved with elastic boundary conditions at *x* = ± 1.8 *nm* in order to set the allowed movement of the particle to 3.6 nm. The elastic boundary condition was imposed by setting the particle flux equal to zero at the boundaries:

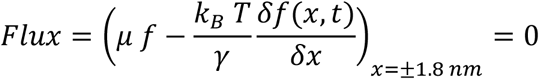

The electrical voltage profile *V*(*x*) was assessed from the net charge density profile *ρ*(*x*), using the following Poisson’s equation

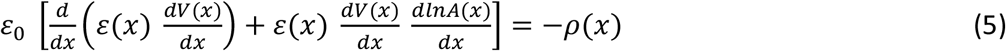

Where *ε*_0_= 8.854·10^-12^ F·m^-1^ is the vacuum permittivity, *ε*(*x*) is the position dependent dielectric coefficient, *V*(*x*) is the electric potential and *A*(*x*) is the position dependent surface. Programs were written in C and are available upon request. Figures were prepared using Origin v4.0 and CorelDraw v18.1.

### Assessment of the energetic profile associated to the voltage sensor position

The energy profiles are assessed self-consistently, by computing (and not assuming) electrostatics. More specifically for each point of the reaction coordinate, i.e. for each position of the voltage sensor along its activation pathway, the corresponding associated energy was assessed by solving iteratively the flux conservative equation for the ionic distribution in the baths and vestibules, and the Poisson’s equation for the voltage profile, until reaching a steady-state (or equilibrium, for equal bath ion concentrations) solution. Once steady-steady (or equilibrium) is reached, the resulting voltage profile is used to assess the electrostatic energy, by integrating the (charge on the voltage sensor)*voltage over the entire space domain (cf the mathematical relationship in the legends of Figures 5 and 7; notice that Figure 7 reports the sole electrostatic energy, while Figure 5 reports the total energy experienced by the voltage sensor, including the mechanical energy due to the attached spring). Obviously the calculation of the energy profile as a function of the voltage sensor position does not involve the solution of the Langevin (or the Fokker-Planck) equation, since in our model these equations are used to assess the movement of the voltage sensor (for example during the evolution of the gating currents), and because we choose the position of the voltage sensor while assessing the energetic profile, as it represents the independent variable of the plot. Thus the electrostatic energy profile shown in Figure 7 essentially comes from the solution of the PNP equations, while varying the position of the voltage sensor. It is important to note that the reliable output of our calculations is the shape of the gating current, and other associated variables. The potential profiles are computed to make our results more understandable and to relate to traditional views of channels. The variable nature of the potential profile is NOT included in this classical view, or in our calculation of potential profiles. Much of the variable nature of the potential profile is included in our solution of the Langevin equations and gating currents, etc.

In our model we assume ionic steady-state equilibrium, i.e., the overdamped system established mathematically by for example (Eisenberg, Kłosek, and Schuss 1995), derived rather than assumed the overdamped high friction limit.This assumption was also used in (Horng et al. 2019), justified by the fact that the voltage sensor moves much more slowly than the ionic re-equilibration in the baths and vestibules. Under this assumption, the energetic profile shown in Figure 5 (i.e. the total energy: electrostatic+mechanical) is exactly the energy encountered by the voltage sensor during its movement, and the energy used to drive the voltage sensor movement through the Langevin (or Fokker-Planck) equation. Mathematically speaking, the spatial gradient of the same energy profile represents the force acting on the voltage sensor, Fex, present in eq. (3) (the Langevin equation) and eqn (4) (the Fokker-Planck equation).

### Assessment of the macroscopic gating current

Once the energetic profile associated with the voltage sensor as a function of its position has been determined (see above), the dynamics (position vs time) of the voltage sensor may be found by solving the Fokker-Planck equation (eqn. 4, with Fex determined by the gradient of the energetic profile). The Fokker-Planck equation gives the time evolution of the probability of finding the voltage sensor in the different allowed positions, assumed that it behaves as a Brownian particle (i.e. its single particle dynamics is described by the Langevin equation 3). Thus at every successive time steps t and t+dt we exactly know (from the numerical solution of the Fokker-Planck equation) how the probability density function describing the voltage sensor position changes (say from f(x,t) to f(x,t+dt)). In response to this voltage sensor movement, also the ions (cation and anion) distributions in the water accessible regions (baths and vestibules) will change (these distributions are assessed together with the energetic profile by solving the PNP equations, as described above), and as a consequence also the total net charge in these regions will change (from Q(t) to Q(t+dt)). For the principle of charge conservation, the net charge difference Q(t+dt)-Q(t) must necessarily come from the left (or right) electrode, and thus the current at the electrode can be derived as (Q(t+dt)-Q(t))/dt. Of course this current will be exactly the same when measured in the two water accessible regions (left and right, as we constantly verified). This method can also be used to assess the current in whatever position along the monodimensional coordinate, by simply assessing the charge accumulated from the edge of the channel up to the position considered, as we did to demonstrate that the total current is conserved in space. Mathematically speaking, the gating current has been determined using the equation:

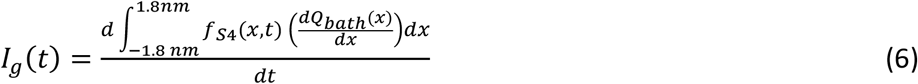

Where 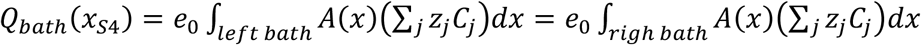 represents the net charge found in the left (or right) bath when the position of the S4 segment is x_S4_, and the integration covers the whole left (right) bath. We checked that the total current density was independent of location x at all times even though the components of the current vary a great deal with x (Eisenberg 2019, 2020; Horng et al. 2019) and of course time *t*.

We emphasize that the main outputs of this paper are the actual calculations of charge movements, potential fields, effective dielectric constants, and most importantly the functional output of the system, the gating currents themselves. The potential profiles used to provide a comforting link with past in Fig. 8 and 9 are most helpful in a human sense, but are likely to be misleading when the more complex interactions described in the last paragraphs are important.

#### Supplementary information

two supplementary Figures, showing an extended sequence alignment for Na_v_ and K_v_ channels voltage sensor and additional information of the water accessibility of the VSD in the I287T Shaker channel accompany this paper.

## Results

### The microscopic scale description

MD simulations performed on available crystal structures can be used to predict the polarizability of a protein or part of it, and thus its effective dielectric constant (Guest, Cashman, and Plotkin 2011; King, Lee, and Warshel 1991; Patargias, Harris, and Harding 2010; Simonson and Perahia 1996; Smith et al. 1993; Voges and Karshikoff 1998; Warshel et al. 2006), a parameter needed in the higher-scale Brownian modeling of voltage gating because it determines the amount of polarization charge available to balance movements of permanent charges, like those of the gating particle. We thus used MD to assess whether the substitution of threonine (found in the Nav channel) in place of isoleucine (found in the Kv channel) at position 287 can sensibly change the polarizability, and consequently the dielectric constant ε of the gating pore, and ultimately the movement and stability of the gating charges inside it.

#### Water accessibility estimates with MD

As a first step we determined the water accessibility of the VSD in the voltage-gated Shaker K channel by performing a MD simulation using the activated state of the Shaker VSD model. After inserting the protein in a phospholipid bilayer and in a water box, and performing the minimization, annealing, and equilibration of the overall system, we ran a 100 ns MD simulation to look for water inside the vestibules and the gating pore. At the end of the simulation we looked at the position of water molecules inside the VSD, to determine whether water could also access the gating pore. The structure on the left of Figure 2A shows a frame at the end of the simulation, while the one on the right shows the water-accessible space of the VSD obtained by the superposition of the water molecules from 60 consecutive frames (one every 0.1 ns). These pictures show that water readily enters both intracellular and extracellular vestibules. The pictures also identify two distinct regions inside the gating pore with different water accessibility. One region, in direct contact with the intracellular vestibule, is totally inaccessible to water, thus forming an effective water inaccessible (WI) region. This region is very narrow – about 2-3 Å, as judged by the distance between the closest water molecules in the two vestibules (Figure 2D). It is located at the level of the gating pore in adjacency to residues F290 on S2, and V236 and I237 on S1 (Figure 2E). The second region, located between the WI region and the extracellular vestibule, including the other seven hydrophobic residues forming the gating pore (including residue I287, Figure 2E), is water accessible (WA). As shown in Figure 2B, the equilibrated simulation shows three water molecules on average in this region (defined as a cylinder of 5 Å radius and extending from −1 Å to 6 Å with respect to the center of mass of the F290 aromatic ring; cf. inset to Figure 2C). These results show the gating pore has two regions with different accessibility to water: a WI region totally inaccessible to water, and an adjacent WA region containing three water molecules most of the time. Due to the different water accessibility, we expect these two regions to have a quite different effective polarizability, with a relative effective dielectric constant close to 4 (the value typical for the interior of proteins) for the WI region, but sensibly higher for the WA region, since here water dipoles could significantly contribute to the stabilization of charges.

**Figure 2.**
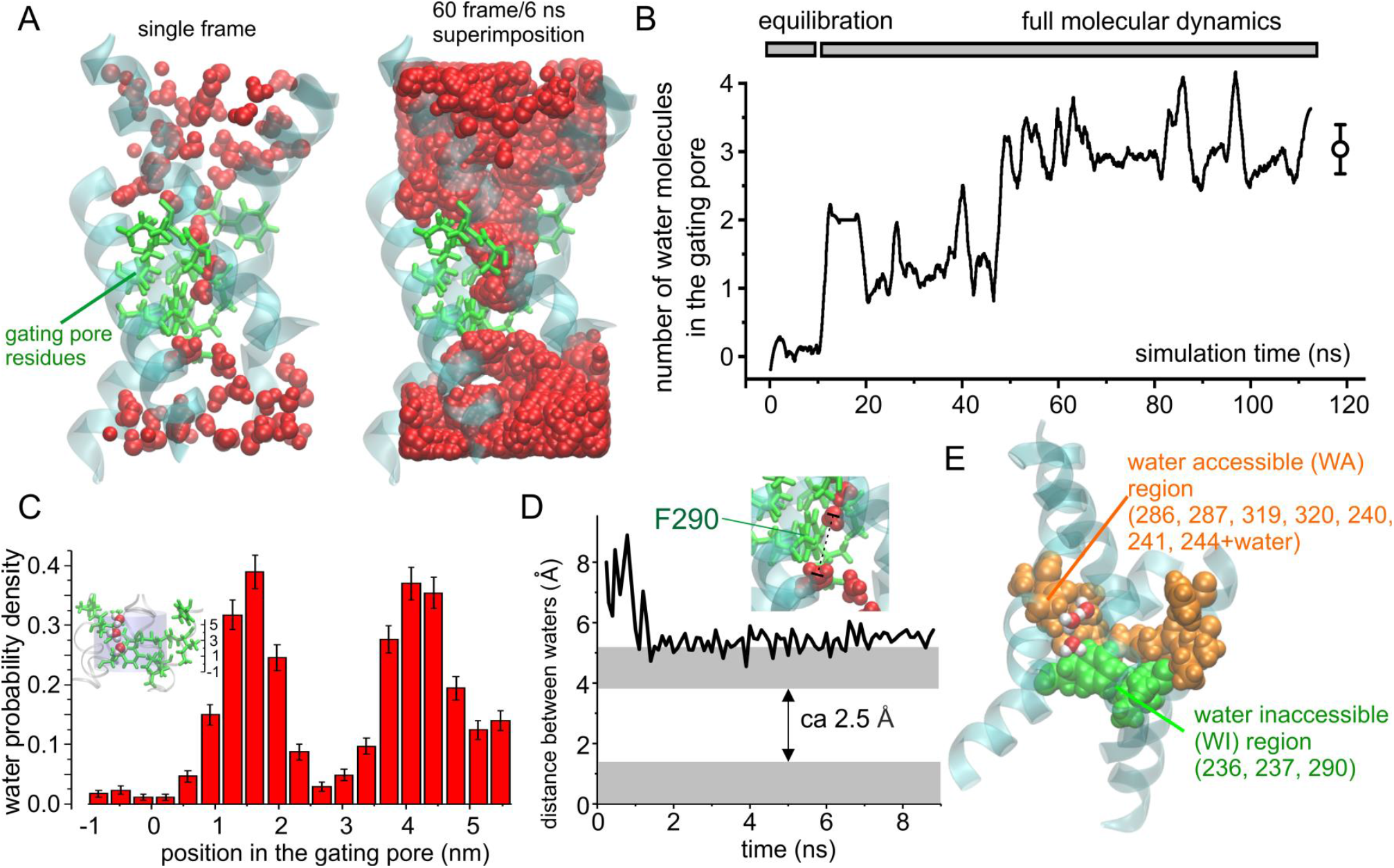
**A) Left:** 3D structure of the Shaker VSD obtained after 100 ns of a MD simulation in which water was allowed to equilibrate inside the intracellular and extracellular vestibules. The protein backbone is represented in cyan/transparent, while the side chains of the gating pore residues are shown in licorice representation, in green. Finally water molecules are in VdW representation, in red. **Right:** Same as the structure shown to the left, but this time the water molecules from 60 consecutive frames of the simulation (one every 0.1 ns) are superimposed, in order to define the water accessible region of the VSD. Notice that only a very short region inside the gating pore is totally inaccessible to water. **B)** Plot of the mean number of water molecules inside the gating pore region (defined as specified in panel C)) as a function of the simulation time. **C)** Plot of the mean number of water molecules in the gating pore. Water molecules were counted when residing inside a cylinder of 5 Å radius and 7Å high, extending from −1Å to +6Å with respect to the z component of the center of mass of the F290 aromatic ring; see inset). **D)** Plot of the distance between the centers of mass of the two closest intracellular and extracellular water molecules as a function of the simulation time. If the radius of a water molecule is 1.4Å (gray regions), a water inaccessible region of 2.5Å is estimated. **E)** 3D structure of the Shaker VSD obtained after 20 ns MD simulation in which water was allowed to equilibrate inside the intracellular and extracellular vestibules. The protein backbone is represented in cyan/transparent, while the side chains of the gating pore residues are shown in VdW representation, in green for the three residues forming the region totally water inaccessible (WI), and orange for the remaining seven residues forming the remaining part of the gating pore, that is partially water accessible (WA). Two water molecules inside the gating pore are also shown (red oxygens and white hydrogens).

#### Estimating the dielectric constant of WA and WI regions of the gating pore using MD simulations

The dielectric constant has two main components, one given by the polarization of the electronic clouds, *ε_el_*, and the other due to the reorientation of permanent dipoles along the applied electric field, *ε_pd_*. Of these, *ε_el_* can be measured experimentally (in bulk solutions) by impedance spectroscopy (Barsoukov and Macdonald 2018; Macdonald 1992) or by the refractive index of the material (Gussoni, Rui, and Zerbi 1998), which depends strictly on the dielectric constant at the appropriate frequency. In addition, the *ε_el_* of a protein region may simply be crudely estimated by summing up the individual polarizability levels of the various amino acids present, that can be obtained experimentally (Voges and Karshikoff 1998). There is regrettably no experimentally recognized method to assess *ε_pd_* inside a protein region, although it is known to depend on dipoles reorientation along the electric field. Indeed, the assignment of polarizability in an atomic scale system has serious ambiguities, as described in (Purcell and Morin 2012). For a small spherical volume of material surrounded by an infinite and homogeneous medium, *ε_pd_* can be estimated by running extended MD simulations and analyzing the equilibrium fluctuation of the total dipole moment of the sample volume, by applying the Kirkwood-Frohlich theory of dielectrics (Frohlich 1949; Simonson and Perahia 1996). Unfortunately, the structure we are studying – the voltage sensor – is immersed in a highly inhomogeneous medium (water and protein residues), on which this approach becomes impractical. In this case the dielectric constant in the small sample volume in the protein interior will also depend on the polarizability of the surrounding medium, whose value is neither known, nor can be assumed to be homogeneous. The assumption of homogeneity is crucial because inhomogeneity creates dielectric boundary charges that can easily dominate the properties of a system (Nadler et al. 2004; Nadler, Hollerbach, and Eisenberg 2003). One can hardly imagine a more inhomogeneous environment than that of a gating pore embedded in a dielectric membrane immersed in an ionic solution.

We thus reasoned that under our conditions the dielectric constant of the sample volume could be estimated from MD simulations run at variable applied electric fields. More specifically, using the basic definition of polarization (P) as the sum of the dipole moments (μ) per unit volume (Vol), and its relation to the dielectric constant (*ε_pd_*), usually valid at moderately low electric fields, we can write:

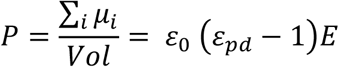

where *ε*_0_ is the permittivity of vacuum, E is the applied electric field, and Vol is the sample volume. The total dipole moment of the sample can be determined using time averaging over the MD simulation (<M>_z_), while applying an electric field along the z direction (E_z_). This allows to calculate ε_pd_ as

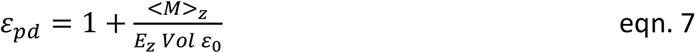

This method emerges naturally from consideration of the relation of charge and electric fields in proteins (Eisenberg, 1996), and has been successfully applied to estimate the dielectric constant of a protein, as well as of water confined inside the pore of a synthetic ion channel (Sansom et al. 1997; Xu, Phillips, and Schulten 1996). In order to validate this method for estimating *ε_pd_*, we first applied it to water molecules in a cubic water box, that is known to have a dielectric constant close to 82 for our TIP3P water model (Simonson 1996). We ran several MD simulations under different electric fields, applied along the z direction, and measured each time the z component of the total dipole moment associated to one water molecule (chosen at random) inside the cubic box. The plot in Figure 3A reports a running average of the total dipole moment assessed for three different applied electric fields, and shows that the value of the dipole moment varies with the applied fields. Figure 3B illustrates the relationship between the total dipole moment along the z direction and the applied electric field. As expected, this relationship is linear for moderate applied fields, while it saturates at higher values. From the slope of this relationship within the linear regime we obtained an effective dielectric constant of about 86, a value quite close to what expected, indicating that our method to estimate the dielectric constant works satisfactorily in this situation. We then applied this method to estimate the effective dielectric constant of the WI and WA regions of the gating pore, both in the WT and in the mutant structure in which isoleucine 287 (found in Kv channels) had been replaced by threonine (mutant I287T), found in Nav channels. In the calculation of the dielectric constant of the WI region we considered all three residues present in this region, namely residues F290, V236 and I237, since they all surround the gating charges while they move through this region of the gating pore (Figure 3D, green residues). As expected for a very hydrophobic, non polar region, the polarizability of the WI region was electronic in nature, with a value around 4.0, irrespective of the residue present at position 287 (Figure 3C, E).

**Figure 3.**
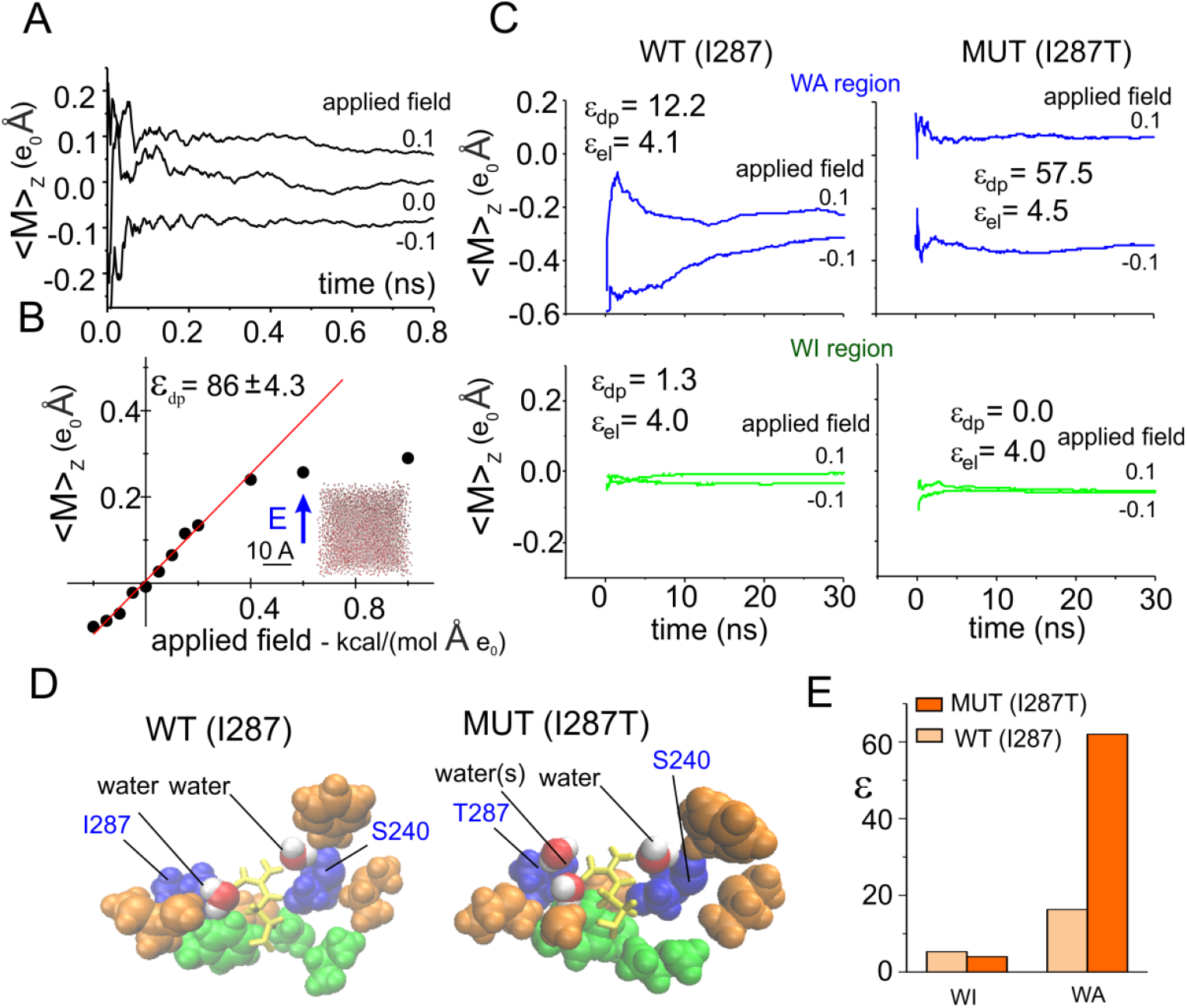
**A)** Running average of the dipole moment assessed for a water molecule inside a 30 Å cubic water box, during three MD simulations performed under different applied electric fields along the z direction. **B)** Plot of the mean total dipole moment of a water molecule as a function of the applied electric field. Each symbol represents the total dipole moment estimated for a water molecule taken at random during the last nanosecond of a 4 ns-long simulations performed in presence of variable applied fields (indicated). Data obtained at applied electric fields of ±0.2 kcal/(mol·Å·e_o_) were fitted with a straight line, and the dielectric constant reported in the plot was estimated from eqn. 7. Notice that 1 kcal/(mol·Å·e_o_) corresponds to about 0.4 Volts/nm. For comparison a constant electric field of 0.2 kcal/(mol·Å·e_o_) would correspond to an electric potential difference of about 240 mV across a phospholipid bylayer. **C)** Plots of the mean total dipole moment at two different applied electric fields, for the water accessible region (top) and for the water inaccessible region (bottom) of the gating pore, assessed from MD simulations using the wild type (WT (I287), left) of the mutated (MUT (I287T), right) structure of the VSD. From these data the ε_pd_ was estimated and reported in each plot. The ε_el_, also reported in the plot, was estimated from the Clausius-Mosotti relationship 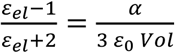, where α is the polarizability and Vol is the volume of the region considered, estimated from the molar polarizability of the amino acidic residues reported in Table 1 of (Voges and Karshikoff 1998). The plot show running averages performed over the last 30 ns simulation, taken from a simulation in wich the indicated electric field was applied for 100 ns to let the molecules to reorient. **D)** 3D structures of the gating pore in the WT and mutated VSD. The gating pore residues are shown in VdW representation, in green or orange/blue depending on their location in the WA or WI region of the gating pore, respectively. Residues 287 and 240, located very close to the gating charge, are colored in blue. Also shown (in VdW representation: red/white) are the water molecules inside the WA region, within 3Å from the charge on R371. The gating charge R371 is shown in licorice representation (yellow). **E)** Plot of the dielectric constant in the WA and WI regions of the gating pore, estimated as *ε* = *ε_el_* + *ε_pd_* as shown in panel C.

The situation is quite different for the WA region where the gating pore widens, and not all residues are close to the moving gating charges. In the assessment of the effective dielectric constant we thus considered only those residues adjacent to the gating charge (i.e. placed at a distance less than 3 Å), namely, residues I287 and S240, plus few water molecules (Figure 3D, blue residues and red/white waters). The dielectric constant estimated in this region was much higher than in the WI region, but, most relevant, it was quite different in the two WA structures considered, being about 20 in the WT and about 60 in the I287T mutant (Figure 3C, E). These results indicate that the nature of the residue at position 287 critically contributes to the polarizability of the WA region around the gating charge, and a mutation of hydrophobic isoleucine at this position introducing a more hydrophilic amino acid like threonine markedly increases the polarization charge in this region.

Two main reasons explain why the WA region has a higher polarizability in the mutant compared to WT. First, the threonine residue has a much higher permanent dipole, because of its hydroxyl group, and thus stronger hydrophilic power than isoleucine. As a result, the dipole moment contributed by this one residue is much higher in the I287T mutant than in the WT structure (Figure 4A). Second, the region close to the gating charge R371 in the I287T mutant appears to have a slightly higher water content than in WT (Figure 4B), presumably because threonine stabilizes nearby water molecules with hydrogen bonds.

**Figure 4.**
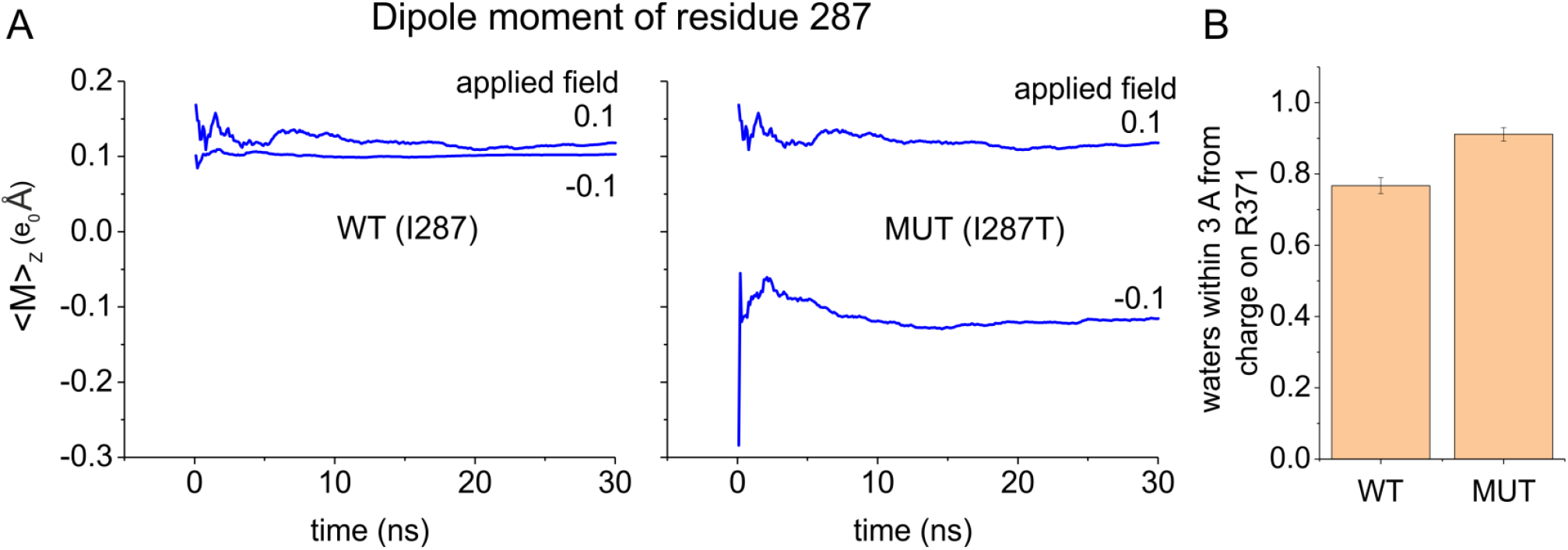
**A)** Running average of the dipole moment associated to the side chain of residue 287 assessed for the WT and the mutated structure, from the same simulations shown in Figure 3C. The mutant threonine is much more polarizable than WT isoleucine. **B)** Mean number of water molecules found inside the WA region of the gating pore and within 3 Å from the charge of R371 in MD simulations performed using either the WT or the I287T mutated structure of the VSD (same simulations shown in panel A).

### The macroscopic scale description

The macroscopic scale description of voltage gating is based on the Brownian model that we have recently developed, and shown to well predict the major signature features of experimental gating currents of Shaker channels (Catacuzzeno and Franciolini 2019; Catacuzzeno et al. 2020) (see Methods). Other papers (Horng et al. 2019; Kim and Warshel 2014; Peyser et al. 2014; Peyser and Nonner 2012b, 2012a) give other views of the voltage sensor and resulting currents at different resolutions. Only (Horng et al. 2019) and (Catacuzzeno and Franciolini 2019) calculate gating currents comparable to those measured experimentally under a range of conditions, with (Horng et al. 2019) having lower atomic detail. Our model takes the relevant structural and electrostatic parameters directly from the 3D structural model of the Shaker VSD (Figure 5A and B). The electro-diffusion of ions in the bath is described through a flux conservation equation that assumes a quasi-equilibrium, on the time scale of gating currents, and the motion of S4 segments is described by Brownian dynamics. The electric potential profile is modeled using the Poisson equation, and includes all the charges present in the system, including the gating charges on the S4 segment and the fixed charges on segments S1-S3 of the VSD. Notably the Poisson equation that is used in our model to assess the electric potential profile contains the relative dielectric constant, ε, as an undetermined parameter that we have estimated above for both the WT and mutant gating pores, using MD simulations. It should clearly be understood that this form of coarse graining has the significant advantage that the lower resolution treatment satisfies conservation laws of mass, charge and current. Coarse graining procedures that do not explicitly introduce the Poisson equation and boundary conditions are unlikely to satisfy conservation laws of charge and current. Periodic boundary conditions in traditional molecular dynamics make it difficult to satisfy Poisson’s equations and conservation of current in a spatially inhomogeneous, non-periodic system like an ion channel protein or gating pore.

**Figure 5.**
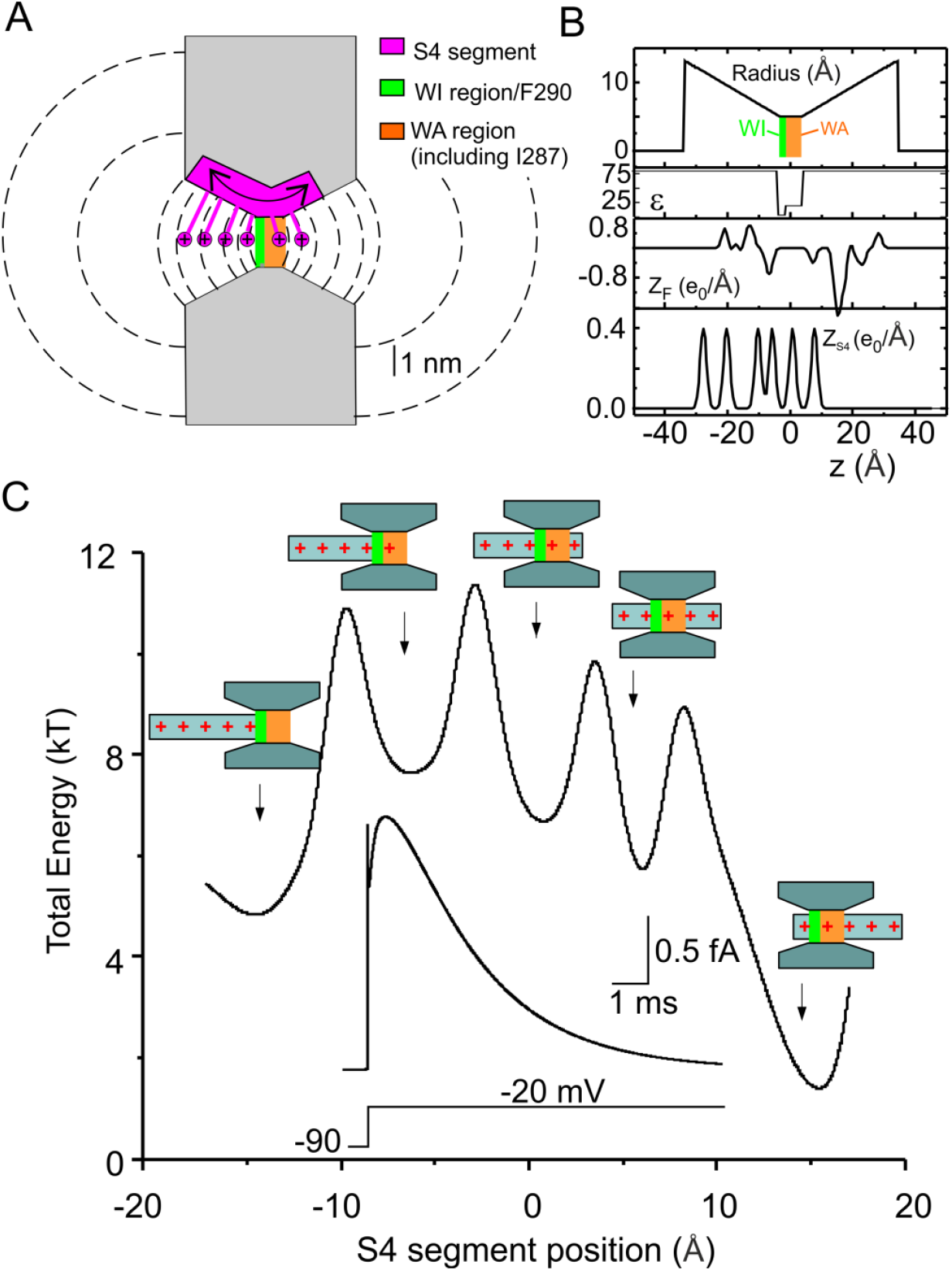
**A)** Schematics showing the geometry of the VSD assumed in our model. The S4 segment containing the gating charges was assumed to move perpendicular to the membrane through the gating pore (7 Å long) and the extracellular and intracellular vestibules (each 31.5 Å long, and opening with a half angle of 15°). Notice that, compared to the model presented in (Catacuzzeno and Franciolini 2019), the gating pore is 5 Å longer to accommodate several hydrophobic residues including residue I287. The vestibules are correspondingly shorter to give the same overall length. The dashed lines represent some of the surfaces delimiting the volume elements considered in our numerical simulations (see (Catacuzzeno and Franciolini 2019) for details). **B)** Profile of the gating pore radius illustrating the 2 Å region, comprising F290 and assumed to be fully impermeant to water (WI, green bar), and the water accessible 5 Å region comprising I287 (WA, red bar), the effective dielectric constant (ε), the fixed charge density located in the S1-S3 region of the VSD (Z_F_), and the charge density on the S4 segment (Z_S4_), for the Shaker model structure. x=0 was placed in the center of the gating pore. **C)** Plot showing the total energy associated to the S4 segment, assessed as 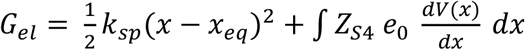, as a function of the S4 segment position. The electrostatic energy profile displays five clear energy wells associated to five different positions of the voltage sensor (represented in schematic form). Inset: Simulated macroscopic gating current evoked by a voltage pulse from −90 to −20 mV, using our Brownian model of the voltage sensor. Note the energy used here does not include entropic effects or frictional losses.

In our previous model we assumed a 2 Å long gating pore localized at the level of F290, that separated electrically the intracellular and extracellular solutions. The effective dielectric constant was set to 4 in this short region, and to 80 elsewhere, including the adjacent region containing residue I287 (Catacuzzeno and Franciolini 2019). The gating pore used in the present study has been changed to account for our MD results shown above. More specifically, the gating pore is now pictured as made of two distinct regions: one with all the properties previously described, with 2 Å length comprising F290, and fully impermeant to water (green bar in Figure 5A and B). The second region, 5 Å long, comprising several hydrophobic residues, including I287, is accessible to water, and has now been assigned a dielectric constant of 20, in accordance with the MD results (Figure 5A and B). As shown in Figure 5C the updated model continues to predict the major features of experimental gating currents obtained in response to a depolarizing pulse. First, a very fast gating current component is present at the beginning of the depolarizing step, rising instantaneously and then falling very rapidly. Second, a later slower current component develops, starting with a plateau/rising phase and proceeding with a slow decay. Both these features have been observed in experiments (Bezanilla 2000; Sigg, Bezanilla, and Stefani 2003). The plot in Figure 5C represents the total energy profile assessed for the various allowed positions of the voltage sensor. It can be seen that the model predicts five energy wells corresponding to the five different positions the S4 segment can assume, in accordance with many studies showing that the voltage sensor moves in multiple steps during the activation process (Bezanilla, Perozo, and Stefani 1994; Henrion et al. 2012; McCormack, Joiner, and Heinemann 1994; Schoppa et al. 1992; Tytgat and Hess 1992; Zagotta, Hoshi, and Aldrich 1994). In our model each step represents the passage of a different gating charge through the high resistance and water inaccessible region of the gating pore (cf. drawings in Figure 5C).

#### Testing the effects of a dielectric constant change on the gating current kinetics

We verified that a different dielectric constant of the WA region can explain the different kinetics of the macroscopic gating current of WT and I287T mutant found experimentally (Lacroix et al. 2013). We used our macroscopic Brownian model of voltage gating to see whether changing the dielectric constant of the WA region from 20 to 60 reproduces the increased rate of gating current seen in mutagenesis experiments.

As shown in Figure 6B and C there was a significant increase in the rate of change of the gating currents evoked by depolarizing pulses, especially at intermediate levels of depolarization (Figure 6C), as observed experimentally. The model also predicted an increased rate of the OFF gating currents induced by repolarization after a short depolarizing pulse (Figure 6E and F). No change was instead observed in the charge vs voltage relationship (Figure 5D). Overall, these results are very similar to those observed experimentally with the mutation I287T, suggesting that a change in the dielectric constant within the gating pore is a major determinant of the functional effects induced by the mutation.

**Figure 6.**
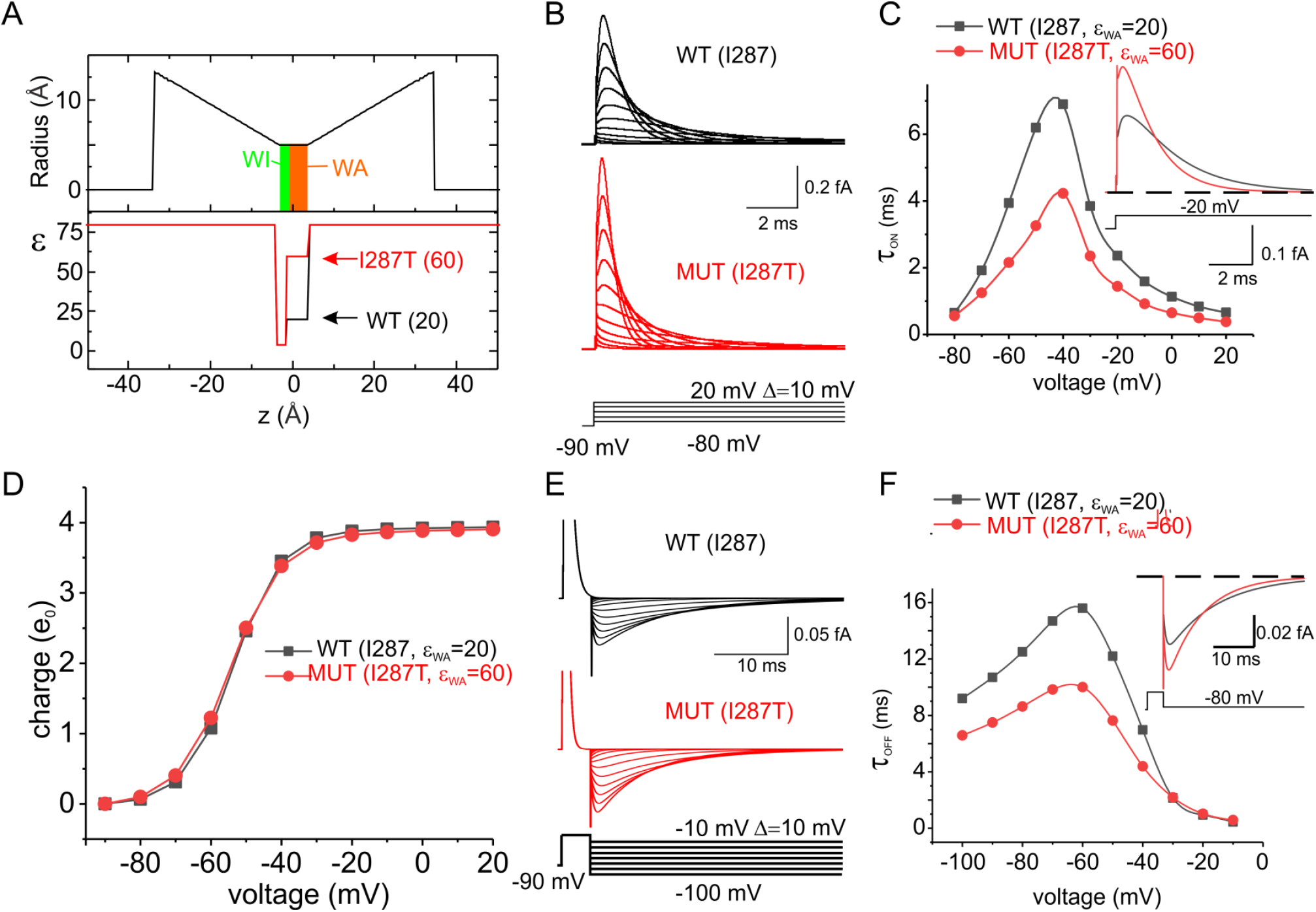
**A)** Profile of the gating pore radius and the effective dielectric constant along the VSD of the WT and the I287T mutated models of voltage gating. The two models only differ in the dielectric constant in the WA region of the gating pore. **B)** Simulated gating currents obtained in response to depolarizing pulses from a holding potential of −90 mV for both the WT (black) and the mutated (red) models. **C)** Plot of the decay time constant of the ON gating currents shown in panel B as a function of voltage. When the gating current decay was clearly bi-exponential, a mean of the two time constants weighted for the respective areas was reported. **Inset:** superimposed ON gating currents obtained in response to a depolarization to −20 mV for the two models. **D)** Plot of the mean charge vs the applied voltage, obtained by integrating the ON gating currents over 40 ms long simulations at different applied voltages. No appreciable difference was observed in the two models. **E)** Simulated gating currents obtained in response to repolarizing pulses after a short (4 ms) depolarizing step to 20 mV, for the WT (black) and the mutated (red) models. **F)** Plot of the decay time constant of the OFF gating currents shown in panel E as a function of voltage. **Inset:** superimposed OFF gating currents obtained in response to a repolarization to −80 mV for the two models.

We then examined the effects of changing the polarizability of the WA region on the electrostatic energy profile, and found that the mutation selectively lowers the first of the four peaks present in the profile (Figure 7). This result appears quite reasonable since the first energetic peak essentially represents the dielectric boundary energy commonly (but incorrectly^1^) called the Born energy for the addition of one (gating) charge to the WA region of the gating pore (Nadler et al. 2004, 2003), and we expect this quantity to depend on the dielectric constant present there (cf. inset to Figure 7). The remaining energy peaks are instead essentially unmodified as the voltage sensor continues to move. This presumably occurs because the charge residing in the WA region is simply replaced by the next charge of the S4 segment, as the voltage sensor moves in response to an applied voltage, keeping the overall charge content in the WA region constant.

**Figure 7.**
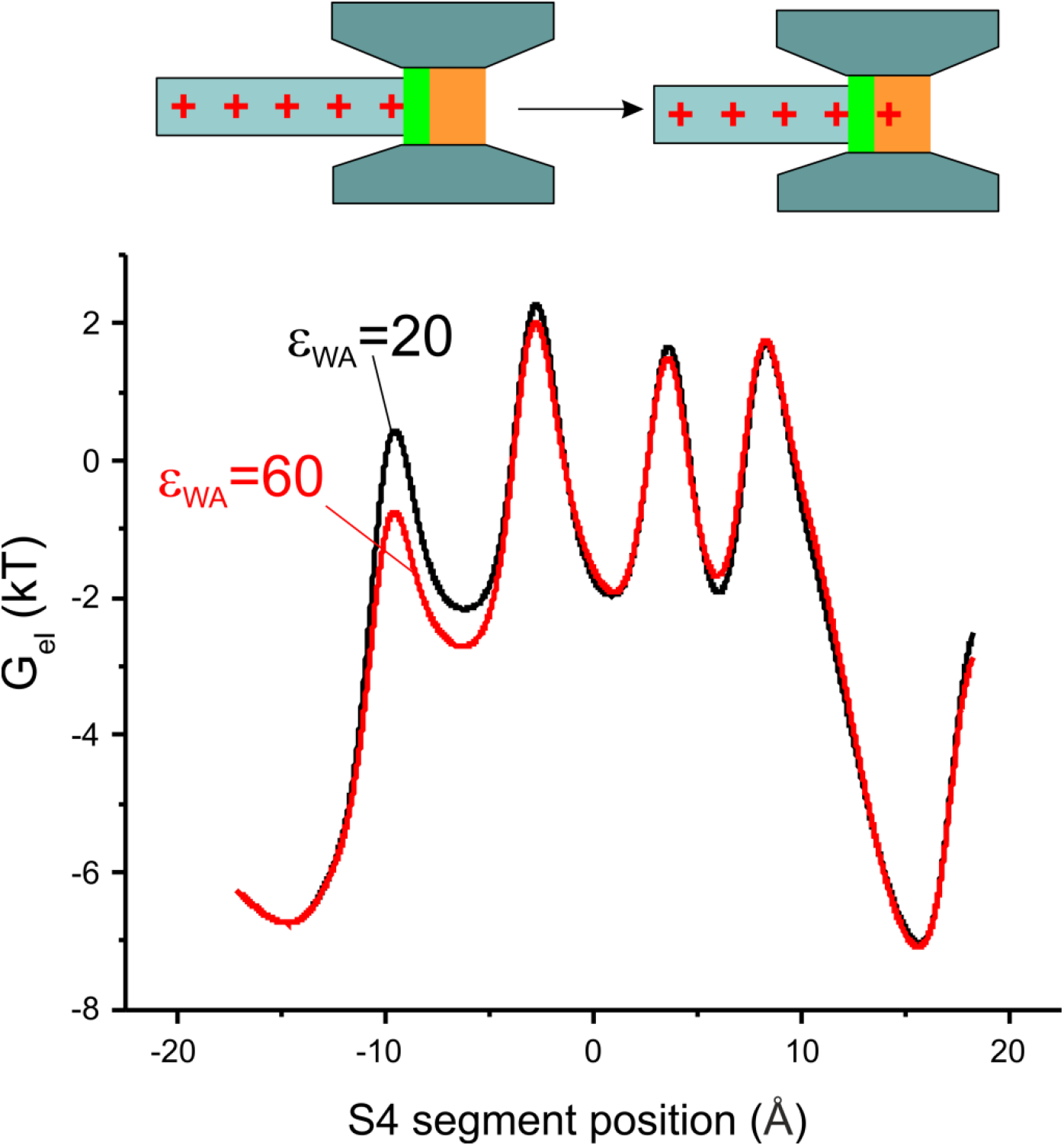
Plot showing the electrostatic energy associated with the S4 segment, assessed as 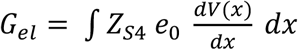, as a function of the S4 segment position, for the two models of voltage-dependent gating differing by the dielectric constant in the WA region of the gating pore. Inset: schematic drawing showing that the change in the electrostatic energy exclusively at the first peak may be explained by the addition of the first gating charge to the WA region of the gating pore.

The energy profile presented in Figure 7 shows that the I287T mutation, beside lowering the first energy barrier reduces also, and by about the same amount, the energy of the neighboring well, corresponding to the first gating charge sitting close to the WA region of the gating pore. If intuitively, a reduction of an energy barrier should lead to a faster activation time course, while a reduction of a local minimum should instead be associated to a slowing of the voltage sensor activation, the overall effect of simultaneous lowering an energy barrier and nearby energy well, is difficult to foresee, and could result in principle in either speeding or slowing the voltage sensor movement, depending on which of the two effects will prevail.

Based on our modeling results we have to conclude that the decrease of the first energy barrier is by far the prevalent effect of the I287T mutation. To verify this conclusion we modeled the effect on gating currents kinetics of altering selectively either the first peak or the second well of the energetic profile, by amounts comparable to those observed following the I287T mutation (Figure 8). The results of these computations show that the effect of altering the energy barrier is much more pronounced than the effect of altering the neighboring local minimum, and thus explain why the mutation results in a net increase in the gating current kinetics.

**Figure 8.**
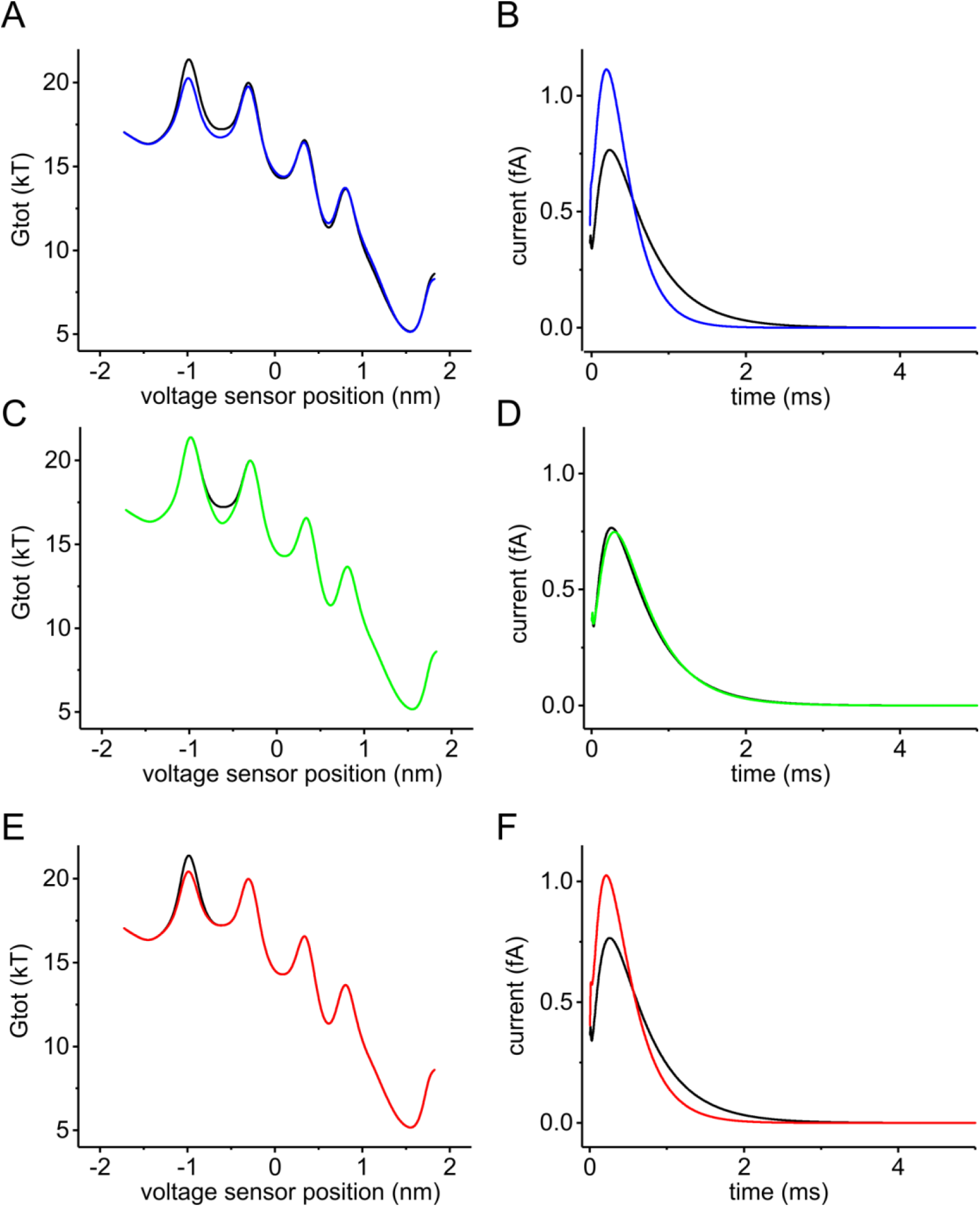
**A)** Energy profiles of the voltage sensor assessed at +30 mV of applied potential using our Brownian model, with a relative dielectric constant of the water accessible region of the gating pore set to either 20 (black) or 60 (blue), in order to predict the effect of the I287T mutation. **B)** Macroscopic gating currents predicted for the energy profiles in A, using the Fokker-Planck equation. The initial condition of the simulation is a probability density function placing all voltage sensors at the first well of the energy profile (x_S4_=-1.425 nm). **C-F)** Starting from the WT model, the energy profile was manually altered to selectively lower the second well (brown trace in C) or the first barrier (green trace in E). The effects of these changes on the time course of the macroscopic gating currents, predicted using the Fokker-Planck equation, are shown in panels D and F, respectively.

## Discussion

Several conclusions may be drawn from the results of our multi-scale analysis. First, using MD simulations we have shown that the gating pore of the Shaker channel, defined in terms of the ten residues proposed by (Lacroix et al. 2014), does not have uniform accessibility to water. More specifically, we have found that the region totally inaccessible to water is actually very short, having a length of 2-2.5 Å and including the three most intracellular residues, namely F290 on the S2 segment, and V236 and I237 on S1. The remaining part of the gating pore, covering a length of about 5 Å, is by contrast accessible to water, with three water molecules often found in this region, as shown by our MD simulations. These results appear in line with previous MD simulations performed on the VSD of other voltage gated ion channels. More specifically, an almost continuous arrangement of water molecules was found in the VSD of the KvAP channel, with a very short interruption at the level of the R133-D62 salt bridge (Freites, Tobias, and White 2006). A similar conclusion was reached with MD simulations on the Shaker homolog Kv1.2, where a very short constriction, localized at the K5-D259 salt bridge, and forming the so-called gating charge transfer center together with F233 (the homolog of F290 in Shaker), prevented communication between the upper and lower crevices (Delemotte et al. 2010; Tao et al. 2010). Finally, a short 2 Å water inaccessible region centered on a conserved phenylalanine was also found in the VSDs of the first three domains of voltage-gated Na channels, and thought to be functionally similar to the VSD of the Kv channel (Gosselin-Badaroudine et al. 2012). The potential difference applied to the membrane will then drop mostly over the very short WI region, because this part of the gating pore is the only region with very high resistance to water and ions. The notion of a very narrow high-resistance region that focuses virtually the entire transmembrane voltage drop has already been suggested by several studies (Ahern and Horn 2005; Asamoah et al. 2003; Islas and Sigworth 2001). The focusing of the electric field is of course incompatible with the idea of ‘constant fields’ that has been a mainstay of biophysics for more than fifty years, and remains so in teaching materials to this day. Electric fields change shape as conditions change because of the fundamental properties of the electric field, as pointed out in the context of channels some years ago (R S Eisenberg 1996; R. S. Eisenberg 1996). F290 has been proposed to act as a gating charge transfer center (Tao et al. 2010), although its role has been later refined to stabilizing the activated VSD conformation (Lacroix et al. 2014; Lacroix and Bezanilla 2011). In our view this residue is important for creating the only very high resistance region inside the VSD, where most of the voltage drops. It is in the 2 Å wide region that includes F290, that the moving gating charges experience the electrostatic force that stabilizes either the closed or open conformation, depending on the applied voltage.

A second result of this study is the quantitative demonstration that the polarizability (i.e., the hydrophobicity) of the residue at position 287 of Shaker channels can greatly modulate the field strength acting on the gating charge as it enters the gating pore. We showed this by estimating the dielectric constant of the region surrounding the gating charge, using MD simulations at variable applied electric fields, and assessing the change in the total dipole moment of the region. In contrast to the computational methods usually employed to assess the dielectric constant (Guest et al. 2011; King et al. 1991; Simonson and Perahia 1996; Smith et al. 1993; Warshel et al. 2006), that are based on the Kirkwood-Frohlich theory of spatially uniform dielectrics (Frohlich 1949; Simonson and Perahia 1996), our method does not assume a homogeneous environment of known polarizability surrounding the sample volume of interest. For this reason our approach can be applied also to restricted regions inside a protein, containing a small number of side chains. This method has been already successfully applied to estimate the dielectric constant of a homogeneous liquid, as well as of water confined inside the pore of a synthetic ion channel (Kolafa and Viererblová 2014; Riniker, Kunz, and Van Gunsteren 2011; Sansom et al. 1997; Xu et al. 1996; Yeh and Berkowitz 1999). However, to the best of our knowledge, it has not been applied before to assess the local dielectric constant inside a (channel) protein.

We found that changing the residue at position 287 significantly changes the effective dielectric constant acting on the gating charge. More specifically, when the isoleucine found at this position in Shaker (as well as in other Kv channels) is changed to threonine, the residue found at the corresponding position in Nav channels, the estimated dielectric constant appears to increase by as much as three times. The observed increase may have two components: i) the much higher permanent dipole present on the threonine side chain (as compared to isoleucine, mostly because of the hydroxyl group there), that contributes directly to increase the polarizability, and ii) the possibility that threonine, as a polarizable dipole, attracts more waters inside the WA region of the gating pore, electrostatically and by establishing weak hydrogen bonds (Perlstein 2001). In summary, our data suggest that the WA region of Nav channel has a higher effective dielectric constant, and this possibly allows the first gating charge to enter the gating pore more easily than in Kv channels. The change in dielectric constant lowers the energetic barrier gating charge encounters in entering the gating pore.

We are aware that our model has limitations. One is related to the estimation of the local dielectric constant inside the gating pore, that we have obtained with classical MD. As there is no electronic polarization with MD, it might be possible that the resulting dipole moment responses are overestimated, and so would be the dipolar component of the dielectric constant. The estimation of the dielectric constant could also be biased by our assumption of a linear relationship between the applied field and the total dipole moment, which we known to be valid in water for moderate field, but do not know to be valid in the complex dielectric environment of the voltage sensor. A second limitation relates to our MD simulations, in two respects. They were run in a limited spatial environment using periodic boundary conditions, and might have produced time varying electric fields as the dimensions of the box change. In addition, their limited time span (100 ns) might not be sufficient to cover wetting/dewetting transitions, which also have an impact on the estimation of the dielectric constant. Nonetheless, in spite of these limitations, we believe that our study provides an important testable hypothesis and opens new avenues for the field of voltage dependent gating.

We also used our Brownian model (Catacuzzeno and Franciolini 2019) to see if the predicted change in the effective dielectric constant found by MD explains the functional effect of the I287T mutation found in experiments (i.e., a noticeable increase in both the ON and OFF gating currents kinetics, but only a small change in the charge-voltage relationship). The output of our Brownian model was quite similar to the experimental results, suggesting that the polarizability change caused by the I287T mutation is a major determinant for the effects observed on the macroscopic gating currents. More specifically we found that the mutation caused a maximum decrease in the time constant of the ON gating currents of ca 57% of its WT value, not very far from the approximately 48% found in experiments (Lacroix et al. 2013). Thus the real effect of the I287T mutation on the gating current kinetics appears slightly higher than that predicted by our multi-scale model. In this regard it should be pointed out that mutagenesis experiments have shown that the size of the residue is also important (Lacroix et al. 2013, 2014). Considering the critical location of this residue, very close to the moving gating charges, it is not surprising that steric effects may also be involved, perhaps by changing the friction experienced by the moving voltage sensor, and thus the rate of the gating currents.

We then looked at the mechanism linking the effective dielectric constant of the WA region to the rate of movement of the voltage sensor by analyzing the effects of the mutation on the energy profile encountered by the voltage sensor as it moves into the gating pore. We found that the reduction in the dielectric constant of the WA region produces a selective decrease in the first of the four energy peaks encountered by the voltage sensor, corresponding to the entry of the first gating charge (R1) into the gating pore. This result is not unexpected since this step involves the entry of a charge into the WA region, a process modulated by its polarizability. It is important to realize that what we show is a prediction of gating currents performed without using adjustable parameters, not a fitting routine where free parameters are adjusted to match the model output with the experimental data (Horng et al. 2019).

In this study we have analyzed in detail the functional properties of Shaker channel gating, with isoleucine or threonine at location 287, since the type of residue present at this position (or equivalent position in Nav channels) is thought to create the kinetic differences between Nav and Kv channels (Lacroix et al. 2013). The results obtained suggest that different activation kinetics of voltage-gated Na and K channels come from different dielectric forces exerted by threonine compared to isoleucine, as the first gating charge enters the WA region of the gating pore. It seems that evolution uses a single side chain to control the speed of the Kv and Nav channels, and ultimately to allow the action potential to exist.

Other processes could contribute to the time course of Na and K channels, as pointed out by Bezanilla’s group. First, residue 363 in S4 is a highly conserved threonine in domains I, II and III of Nav channels, while is a highly conserved isoleucine in Kv channels. Mutation of this isoleucine to threonine makes the voltage sensor of Kv channels move faster, as inferred from the time course of the gating currents (Lacroix et al. 2013). From the position of this residue relatively far from the gating charge R1, we suspect that its contribution to the polarizability would not be significant, suggesting that a different mechanism is likely responsible for this effect. Second, the presence of the β1 subunit of Nav channels also accelerates the voltage sensor, by an unknown mechanism (Lacroix et al. 2013). Finally, the sensors in Nav channels exhibit positive cooperativity which also accelerates the overall speed of channel activation (Chanda, Asamoah, and Bezanilla 2004).

We point out that (within the limitations of our models) what we propose is both logically sufficient and logically necessary to account for the difference in speed and the structural movements of the voltage sensor. It is sufficient because we reproduce the properties of gating current under a realistic range of conditions. It is necessary because the movement of charges that we calculate must produce the currents we calculate given the universal and exact nature of the Maxwell equations that link charge and current. Of course, evolution is not logical. It often provides redundant mechanisms that are beyond the necessary and (perhaps barely) sufficient. These redundant mechanisms may provide properties not glimpsed in gating currents studied with step functions in the way they are usually measured. These mechanisms may make the gating system robust in conditions we have not explored.

A similar multi-scale approach may also be used in the future to understand the functional effects of the many other mutations that have been performed on the VSD, and so to have a physical interpretation of the data and a more advanced understanding of voltage-dependent gating.

## Acknowledgements

This work was supported by Grant Ricerca di Base 2017, from the University of Perugia. We are grateful to Prof. Bezanilla for teaching us the importance of his results in a discussion that motivated the models of this paper. We thank the referees for suggesting significant improvements to our calculations and presentation of results.

## Supplementary Material

**Figure S1.**
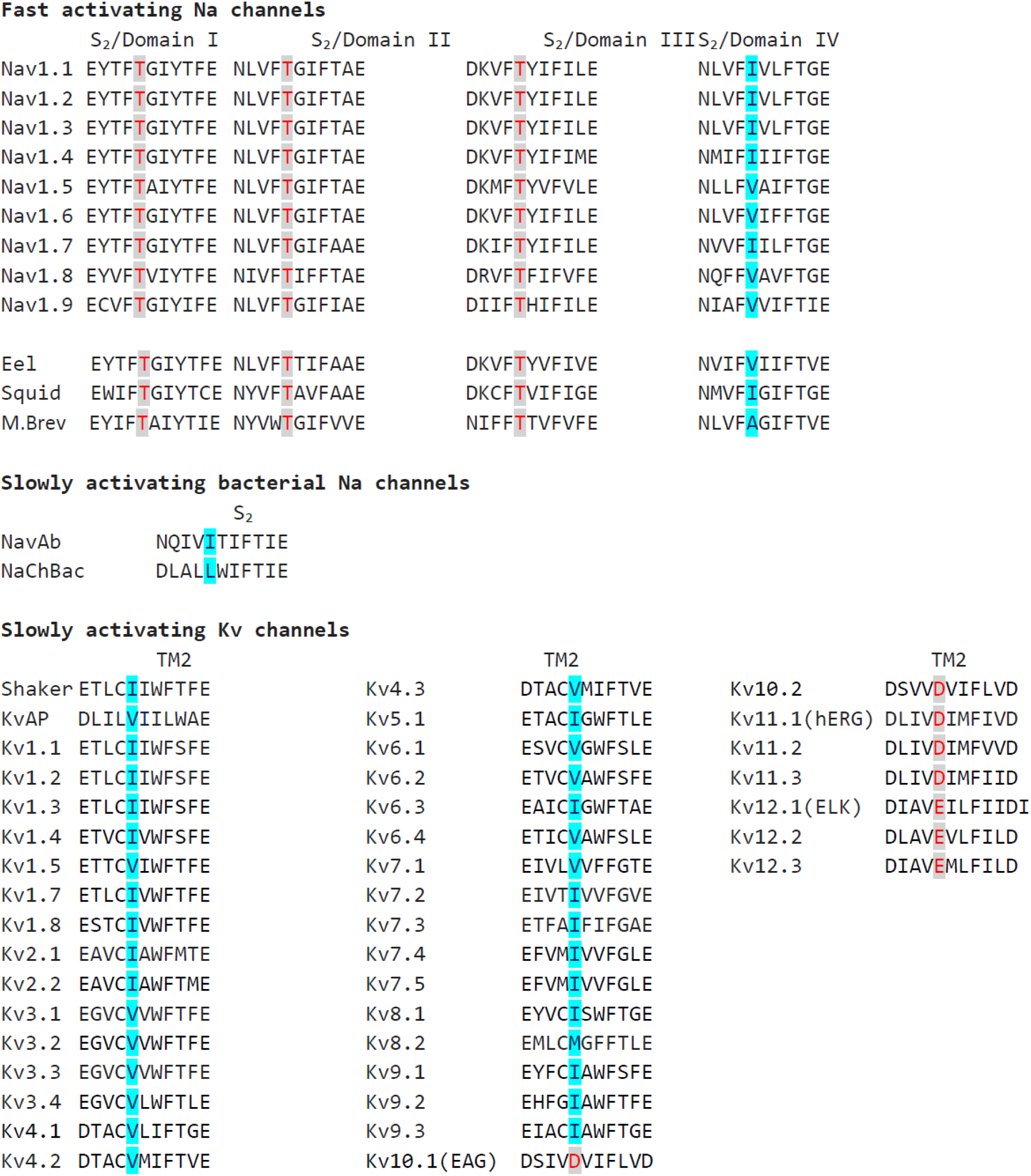
Conservation of the speed-control residue in the S_2_ segment in different types of fast and slowly activating Nav, and slowly activating Kv channels. The residues homologous to Shaker I287 are marked in red if hydrophobic (E, D) and in blue if hydrophilic (I, V, M). The data clearly show that in the first three domains of all the reported fast activating Na channels the position homologous to Shaker I287 is occupied by a threonine residue, confirming the functional importance of the hydrophilic threonine at this position. The IV domain of fast activating Nav channels shows instead either an isoleucine or a Vvaline at the corresponding position, in accordance with a functional role of this domain not primarily involved in the activation, but in the slower inactivation process. As for the slowly activating bacterial Na channels, hydrophobic isoluecine or Leucine is found, again in accordance with a direct correspondence between the hydrophilicity of this residue and the rate of voltage sensor activation. Finally, as in Shaker channels, also in KvAP and in the 31 members of the first 9 families of human Kv channels the position homologous to Shaker I287 is occupied by a hydrophobic residue (Isoleucine o Valine, Methionine in one case), thus strengthening the notion of a strict association between the slow voltage activation of Kv channels and the presence of a hydrophobic residue at this position. This association is instead not respected in the last three families of Kv channels (EAG or Kv10, ERG or Kv11, and ELK or Kv12) where a negatively charged residue at the position homologous to Shaker I287 is always found. Although we have not a clear explanation for this finding, it is interesting to notice that for these groups of Kv channels a different inter-domain assembly, with a non-swapped topology between VSD and pore domain, has been found (Butler et al. 2020; Wang and MacKinnon 2017; Whicher and MacKinnon 2016). It is also interesting to notice that MTSET accessibility data indicate that in hERG Kv channels the extracellular water crevice enters into the VSD well beyond the residue D460 (homologous to Shaker I287) and F463 (homologous to Shaker F290). Thus the organization of the VSD in these channels appears markedly different as compared to the Kv channels with a swapped-domain topology, of interest in our study.

**Figure 2.**
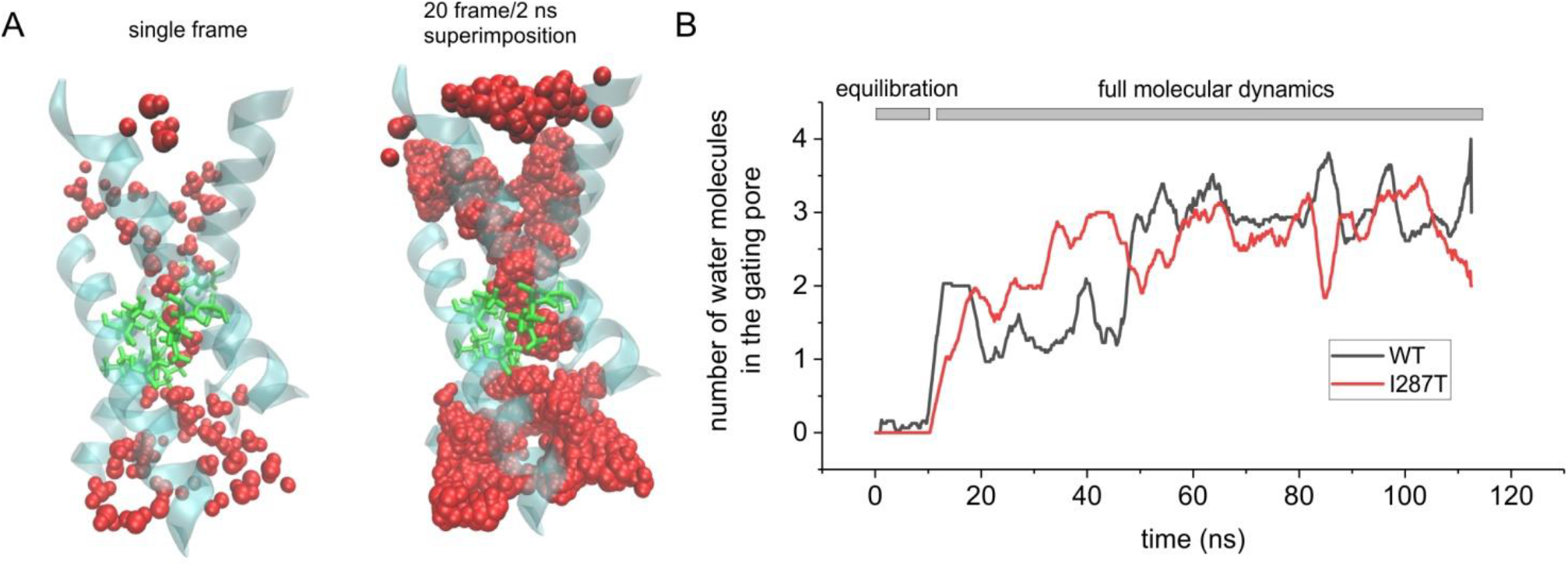
**A) Left:** 3D structure of the Shaker VSD carrying the mutation I287T, obtained after 100 ns of a MD simulation in which water was allowed to equilibrate inside the intracellular and extracellular vestibules. The protein backbone is represented in cyan/transparent, the side chains of the gating pore residues are shown in licorice representation (green), and water molecules in VdW representation (red). **Right:** Same as the structure shown to the left, but here the water molecules from 20 consecutive frames of the simulation (one every 0.1 ns) are superimposed, in order to define the water accessible region of the VSD. Notice that only a very short region inside the gating pore is totally inaccessible to water. **B)** Plot of the mean number of water molecules inside the gating pore region (defined as the water molecules residing inside a cylinder of 5Å radius, extending from −1Å to +6Å with respect to the z component of the center of mass of the F290 aromatic ring) as a function of the simulation time. The black and red lines refer to the simulation performed with the WT structure (the same shown in Figure 2 of the main paper) and with the mutant (I287T) structure, respectively.

## Footnotes

1 The Born energy was computed by Born in an infinite homogeneous domain without dielectric discontinuities. The system we study is about as far from homogeneous as one could imagine and its biological functions are determined by the heterogeneous distribution of polarization and dielectric properties. The use of the phrase Born energy is seriously misleading in the opinion of at least one of the authors.

